# Structure and catalytic mechanism of methylisocitrate lyase, a potential drug target against Coxiella burnetii

**DOI:** 10.1101/2025.01.06.631558

**Authors:** William S. Stuart, Christopher H. Jenkins, Philip M. Ireland, Michail N. Isupov, Isobel H. Norville, Nicholas J. Harmer

## Abstract

We present a comprehensive investigation into the catalytic mechanism of methylisocitrate lyase, a potential drug target candidate against the zoonotic pathogen *Coxiella burnetii*, the causative agent of Q fever and a federal select agent. Current treatment regimens are prolonged, often with incomplete clearance of the pathogen. We utilised a structure-based bioinformatics pipeline to identify methylisocitrate lyase as a candidate therapeutic target against *C. burnetii* from a list of essential genes. Wild-type *C. burnetii* methylisocitrate lyase has a *k_cat_* of 32,000 s^-1^ (compared to 105 s^-1^ for *Salmonella enterica*) and isocitrate inhibits with a *K_I_* of 6 mM. We have determined the previously uncharacterised substrate-bound structure of this enzyme family, alongside product and inhibitor-bound structures. These structures of wild-type enzyme reveal that in the active state the catalytic C118 is positioned 2.98 Å from O5 of methylisocitrate and Arg152 moves towards the substrate relative to the inhibitor bound structure. Analysis of structure-based mutants reveals that Arg152 and Glu110 are both essential for catalysis. We suggest that Arg152 acts as the catalytic base that initiates the methylisocitrate lyase reaction. These results deepen our understanding of the catalytic mechanism of methylisocitrate lyase and could aid the development of new therapeutics against *C. burnetii*.

## Introduction

The gammaproteobacterium *Coxiella burnetii* is an obligate intracellular pathogen that infects a wide range of mammals and other species [4]. *C. burnetii* has a biphasic lifecycle with large and small cell variants (LCV and SCV, respectively). The SCV has a spore-like morphology, enabling airborne spread as the most common route of infection in humans, resulting in the disease Q fever [5, 6]. Previous outbreaks have caused significant societal and economic damage through human disease and culling of infected livestock. The three year outbreak in the Netherlands between 2007 and 2010 is estimated to have cost up to 600 million Euros [7]. Q fever initially presents as an acute phase characterised by fever-like symptoms which resolve without treatment in most cases. Approximately 10% of symptomatic *C. burnetii* infections develop into chronic illness that can lead to Q fever fatigue syndrome (QFS) [8, 9]. *C. burnetii* is an intracellular pathogen that subverts the lysosomal pathway to establish a unique replicative vesicle in the host cell, the *Coxiella* containing vacuole (CCV) [10]. Within the pH ∼5.0 CCV, *C. burnetii* is shielded from the host immune system, can obtain essential nutrients from the host and is able to secrete effector proteins to manipulate host cell function [11]. The bacterium has a reduced genome with around 2100 coding sequences [12] and requires a supply of most amino acids from the host. *C. burnetii* establishes a bipartite metabolism with the host cell (a feature shared with the closely related pathogen *Legionella pneumophila*), preferentially using host amino acids for carbon and energy [13]. As *C. burnetii* is highly auxotrophic, laboratory cultivation requires a defined axenic media comprised of cell culture media, peptone, and a complex mixture of additives [14].

Eradication of *C. burnetii* in chronic Q fever cases is highly challenging due to the slow growth rate of *Coxiella* and its intracellular location [10]. The current gold standard therapeutic strategy consists of a combination of doxycycline and hydroxychloroquine [15]. This treatment regimen typically lasts for at least 18 months and consists of three doses of doxycycline and two doses of hydroxychloroquine daily [16]. Doxycycline particularly is associated with significant side effects, which may lead to poor patient adherence [17].

Given the challenges of the current therapeutic strategy there is a need for new drugs targeting *C. burnetii*.

Comparative genomics of *C. burnetii* revealed a “core” genome of approximately 1300 genes that is conserved across a wide range of pathogenic and environmental isolates [18]. 466 genes were identified as essential for survival in culture using a transposon mutagenesis screen [19], the large majority from this core genome. Many of these genes are essential in other bacteria, but some are likely more specific to *Coxiella*’s unique lifestyle. By *in silico* screening of the essential genes, we identified methylisocitrate lyase (PrpB) as a promising candidate.

PrpB catalyses the lysis of 2-methylisocitrate (2-MIC) into succinate and pyruvate and is the final step in the methylcitrate cycle [20]. The methylcitrate cycle functions in bacteria to process propionate, an intermediate in catabolism of several amino acids (particularly valine), into central metabolism (Figure 1a) [21, 22]. Accumulation of propionate is toxic to most bacteria [23]. The related enzyme isocitrate lyase (ICL) (a putative drug target in *Mycobacterium tuberculosis* and other pathogens) is the first step in the “glyoxylate shunt” that acts as an alternative to one half of the citric acid cycle [24, 25]. ICL has activity on 2- MIC, enabling the clearance of propionate independent of PrpB [26]. However, *C. burnetii* lacks a gene for ICL (and the remaining glyoxylate shunt enzymes), potentially explaining why PrpB is essential for survival of this pathogen.

**Figure 1:**
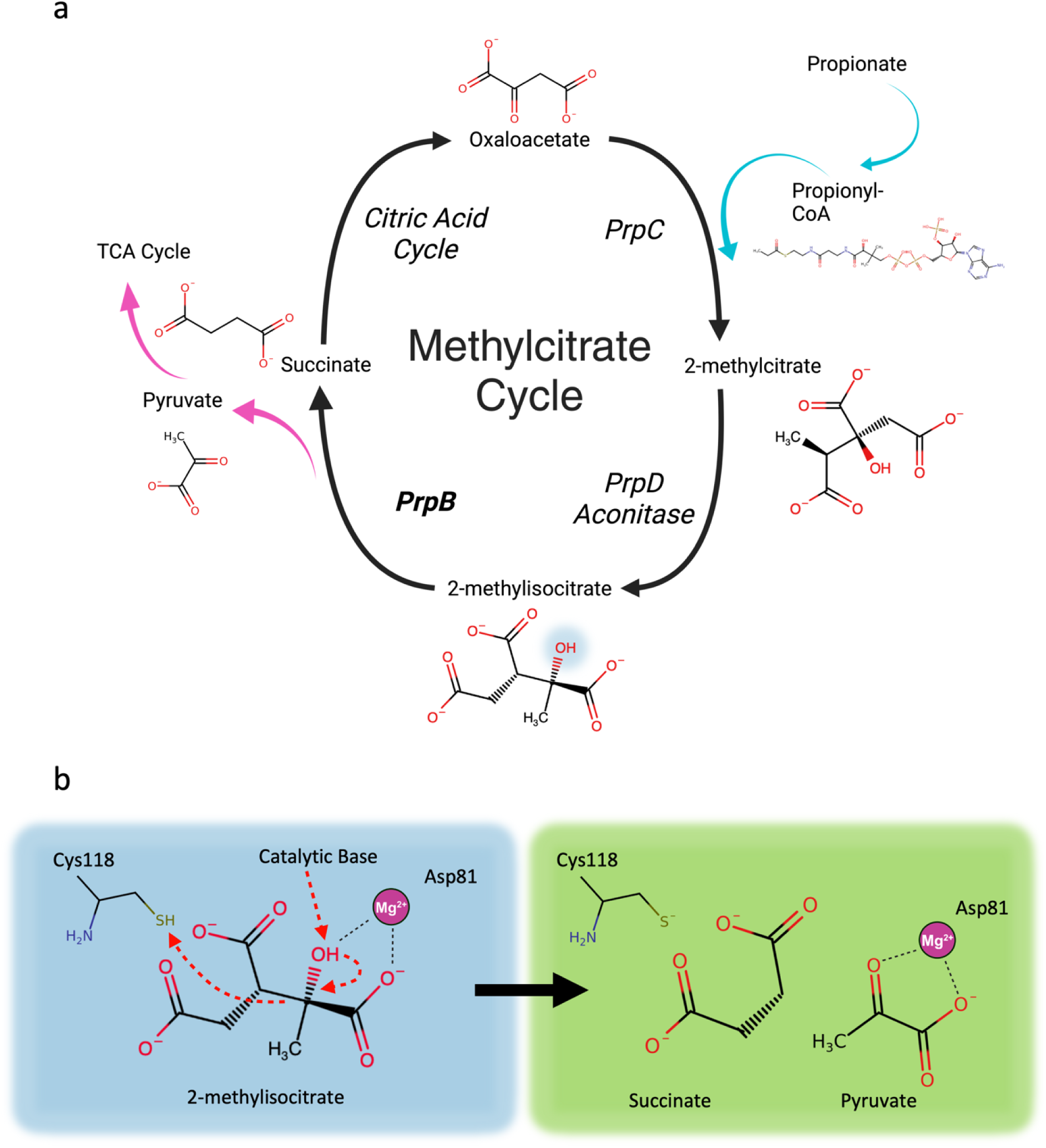
Role and function of 2-methylisocitrate lyase. **a**: Overview of the methylcitrate cycle showing the clearance of the toxic metabolite propionate. The highlighted 2-methylisocitrate hydroxyl group (bottom of image) is shifted by one carbon compared to 2-methylcitrate which is necessary to give lysis products that feed into the TCA cycle. **b**: Reaction mechanism of 2-methylisocitrate lyase. Left hand side shows the substrate and reaction progression with proton abstraction by an unidentified base and proton donation from cysteine 118. Right hand shows products succinate and pyruvate following cleavage of C2-C3 bond. Panels a and b created in BioRender.com and Microsoft PowerPoint.

PrpB and ICL are homo-tetrameric enzymes with a molecular weight approximately 140 kDa [27]. Each protomer contains one active site that is organised around a bound magnesium ion that coordinates the substrate [28]. After substrate binding, a mobile loop changes conformation to close over the active site, sequestering solvent and bringing the catalytic residues into proximity [29]. The catalytic mechanisms proposed for ICL and PrpB are very similar, with a catalytic base deprotonating the substrate hydroxyl group to break a carbon-carbon bond; following this, a catalytic acid protonates the four carbon intermediate to form succinate (Figure 1b). Cysteine 118 (*C. burnetii* PrpB numbering), located on the mobile loop, is proposed to be the catalytic acid (proton donor) that completes the lysis reaction [30]. However, there remains some controversy as to the identity of the catalytic base with several possible side chains proposed, including Y40, E110 and R152 or a water molecule (deprotonated by D53; *C. burnetii* PrpB numbering) [27, 31]; whilst the equivalents of Y40, H108, and R152 have been proposed for ICL [32, 33].

Here, we present the structure of *C. burnetii* PrpB in its apo form and bound to substrate, product and an inhibitor. This provides the most comprehensive structural study of a methylisocitrate lyase to date and provides a substrate bound structure, previously uncharacterised for any 2-MIC or isocitrate lyase, at 1.9 Å resolution. We have characterised the kinetic parameters of the wild-type enzyme and selected point mutants suggested by the structural results. Together our results enable a clearer insight into the catalytic mechanism of PrpB and suggest that, surprisingly, Arg152 is the catalytic base. Our evidence suggests that PrpB is an attractive target for a new antimicrobial.

## Results

### AI Structure Prediction to Triage Essential Genes for Target Identification

Drug target candidate selection (Figure 2) built upon a previous transposon mutagenesis screen (TraDIS) which identified 466 putative essential genes within the genome of *C. burnetii* Nine Mile Phase II [19]. The first stage in the selection pipeline provided general information on all essential genes. This included extracting protein mass from UniProt [34], batch predicting isoelectric point using IPC [35] and the number of transmembrane domains (TMDs) in each gene using TMHMM v2.0 [36]. We rejected genes that contained two or more predicted TMDs (74/466), along with 16 transposases. BLAST [37] similarities for all essential genes against the human proteome were generated, with 209/466 returning an E-value of less than 0.05, indicating some similarity with a human protein. Since this may lead to off-target effects they were rejected. Together these criteria removed 277 genes (with some rejected on multiple criteria). Each target was also run through BLAST against the PDB [38], returning 50 with an E-value of greater than 0.05. These proteins were also rejected, since without a close homologue in the PDB they may be more difficult to produce and characterise functionally and are likely to have poorer performance in structure prediction. This triaging process reduced the list of possible targets from 466 to 139. BLAST similarities were calculated against the proteomes of eight other pathogens, to select conserved targets, as a broader spectrum target would give greater value to development. *Legionella pneumophila* (the closest pathogenic relative of *C. burnetii*); *Francisella tularensis*, *Yersinia pestis*, *Burkholderia pseudomallei*, and *Bacillus anthracis* are additional Tier 1 select agents [39]; and *Pseudomonas aeruginosa*, *Klebsiella pneumoniae* and *Acinetobacter baumannii* are Gram-negative “ESKAPE” pathogens of the greatest concern for antimicrobial resistance [40].

**Figure 2:**
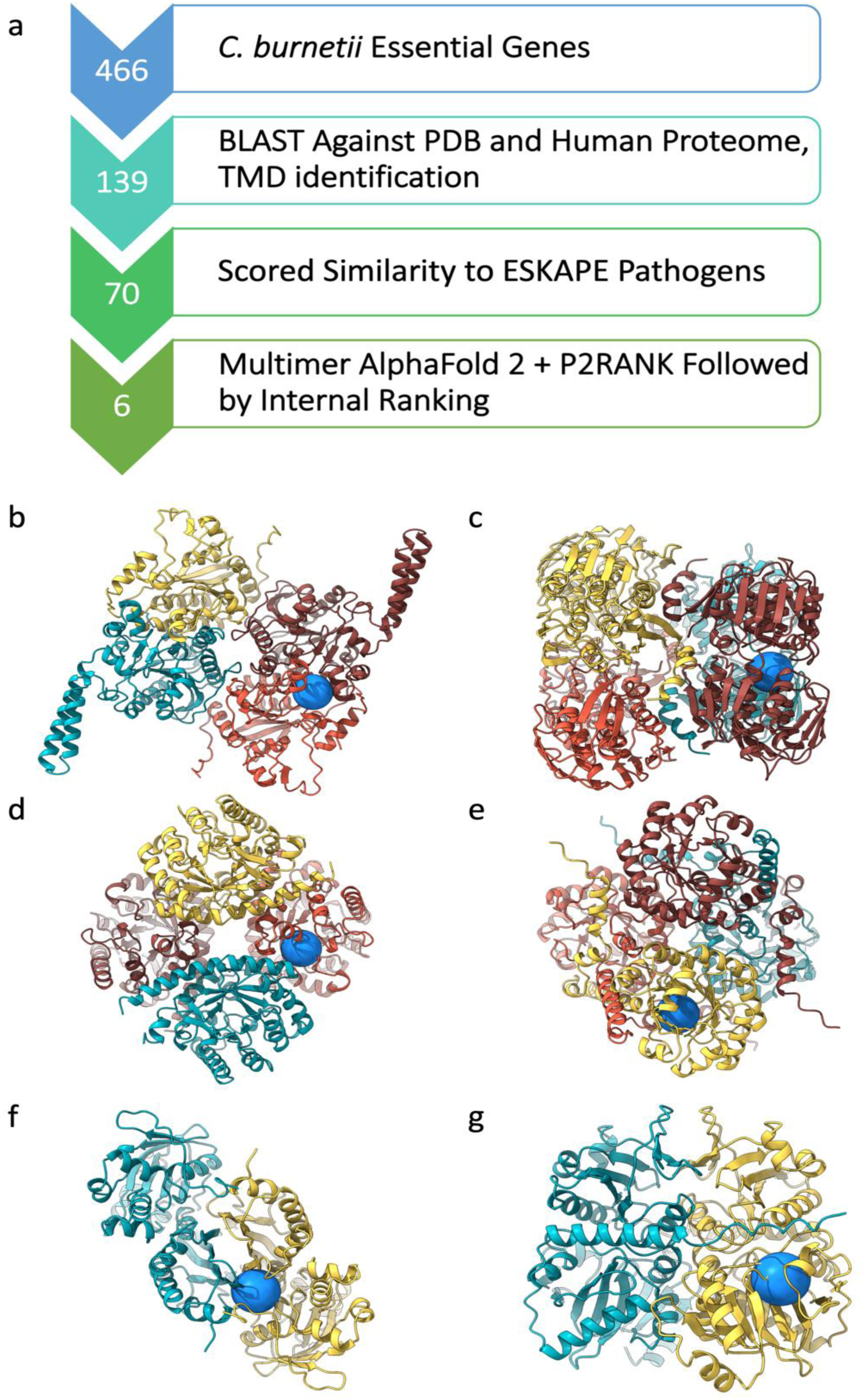
Identification of C. burnetii Drug Targets. **a**: Flow diagram of downselection process for essential genes in C. burnetii, resulting in a small number of candidate targets. **b-g**: AlphaFold predicted structures of C. burnetii drug target complexes from in-house script coloured by chain with highest scoring predicted drug pocket site from P2RANK shown by the blue sphere. **b**: Acetyl-CoA carboxylase (CBU_1510 and CBU_0893). **c**: UDP-N-acetylglucosamine 1- carboxyvinyltransferase (CBU_0751). **d**: Dihydropteroate synthase (CBU_1351). **e**: 2-methylisocitrate lyase (CBU_0771). **f**: Thiamine monophosphate kinase (CBU_1415). **g**: D-alanine D-alanine ligase (CBU_1338). Panels b-g were generated with UCSF ChimeraX 1.7 [3].

We devised the following scoring scheme to rank targets by cross-compatibility with other pathogens. Proteins that had an E-value similarity to *C. burnetii* of greater than 1x10^-20^ received 0, those less than 1x10^-60^ received 3 and those in between a score of 2. Scores against all pathogens were totalled and scaled from 0-100, with a score of 100 indicating a gene had an E-value of less than 1x10^-60^ against all organisms. This resulted in 70 targets having a score of above 70 (Figure 2) while 31 had a score of 100, indicating high conservation across the eight other pathogens.

Targets scoring above 70 were taken forward for structure prediction using AlphaFold v2.1 [41]. Where possible, predictions were made using the predicted biological assembly to ensure the correct construction of the active site. We wrote a script to automate construction of AlphaFold input files to ensure consistency (Supplementary Files). Key inputs are a list of corresponding PDB codes (identified in the target identification spreadsheet) and the target organism. The PDB1 file (which contains the complete biological assembly) is downloaded and the correct sequence FASTA file is generated for the specified organism using BLAST and the biological assembly data. Targets are then predicted using AlphaFold multimer. This script can be used for any organism and therefore may be of value to others. 25 multimeric targets were successfully predicted while four targets failed due to memory limits or incorrect predictions.

Output structures were subsequently passed to pocket prediction analysis using P2RANK [42]. This was carried out on the predicted monomer or multimeric structures. Manual inspection of P2RANK pocket scores indicated that pocket location prediction reflected the location of ligand binding sites where these were known from ortholog structures. In several cases it was seen that utilisation of AlphaFold multimer predicted structures improved the P2RANK pocket score, justifying the approach to incorporate protein complex prediction.

P2RANK highlighted 16 potential candidates. We triaged these by prioritising those likely to have off-target effects (ATP or GTP binding proteins); proteins involved in cell wall or outer membrane biosynthesis, as *C. burnetii* shows little susceptibility to current antibiotics targeting these [16]; and enzymes for which substrates are highly challenging to obtain. This left a final set of six candidates, shown in Figure 2, which were subject to recombinant expression and purification was attempted. Methylisocitrate lyase (PrpB; *CBU_0771*) was readily optimised for robust expression in *E. coli* and enzyme activity. We focus here on a detailed analysis of this target.

### C. burnetii PrpB Kinetic Analysis

We implemented an assay to follow *C. burnetii* PrpB (*Cb*PrpB; Supplementary Figures 1 and 2) lysis of 2-MIC by purified by coupling generation of the product pyruvate to oxidation of nicotinamide adenine dinucleotide (NADH) utilising lactate dehydrogenase [43]. Our results indicate that wild-type *Cb*PrpB has a Michaelis constant (*K_M_*) of 390 µM, approximately one order of magnitude greater than the value reported for *E. coli* (Figure 3). Experiments did not provide strong evidence for cooperativity in the tetrameric enzyme. *Aspergillus fumigatus* PrpB is inhibited by isocitric acid, with a Morrison *K_i_* of 258 µM when tested with 2-MIC at this enzyme’s *K_M_* of 18 µM [44]. In this study, a 2-MIC concentration of 500 µM, gave a Morrison *K_i_* of 6 mM (Figure 3B).

**Figure 3:**
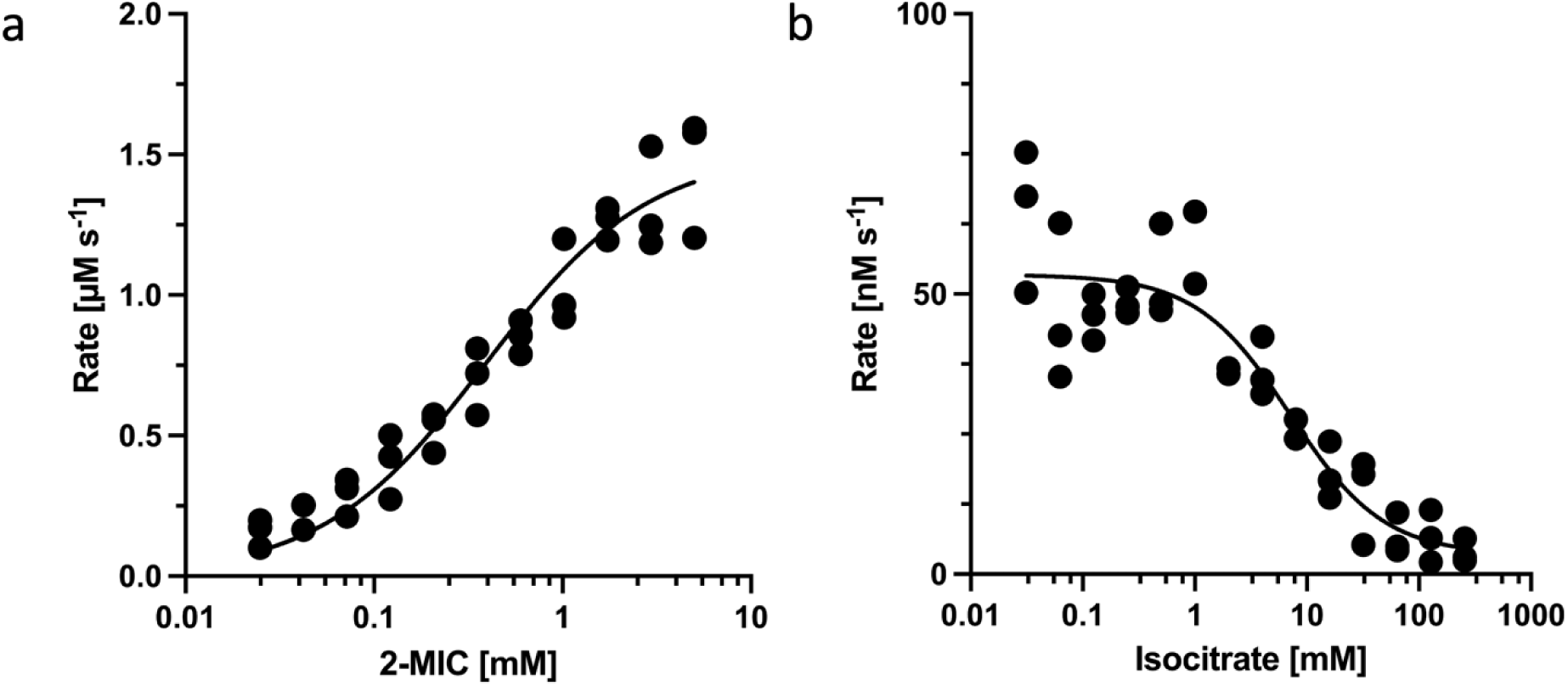
Enzyme kinetics for C. burnetii 2-methylisocitrate lyase. **a**: Michaelis -Menten assay using 50 pM PrpB shows the enzyme has a K_M_ of 390 ± 47 µM. **b**: Morrison K_I_ assay at constant concentrations of 50 pM PrpB and 500 µM 2-MIC gives an inhibition constant K_I_ of 6 ± 2 mM. Kinetic parameters and plots generated with Graphpad Prism (v10.1.1 for MacOS, GraphPad Software, www.graphpad.com).

### Structural Investigations

*Cb*PrpB produces crystals readily in a range of conditions. These were amenable to soaking with substrates, products, and inhibitors. This enabled us to gather the most extensive set of ligand-bound structures obtained for any PrpB to date (Table 1). Six crystal structures were determined in three different space groups, at resolutions of between 1.6 and 2.5 Å. The asymmetric units contained either two, eight or twelve molecules (Supplementary Tables 1 and 2). As with other PrpB and ICL structures, *Cb*PrpB forms a homotetramer.

**Table 1:**
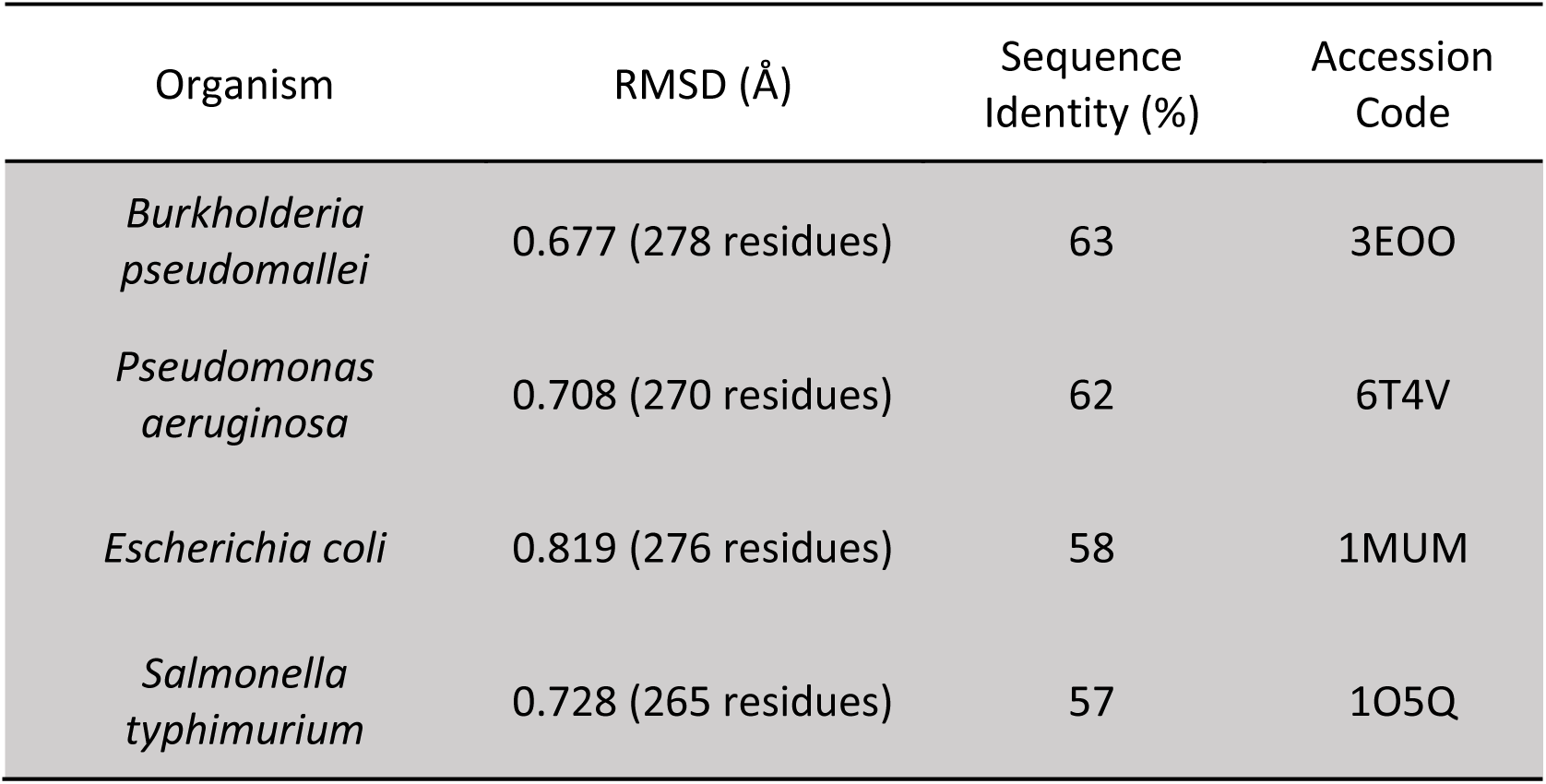
Comparison of apo C. burnetii PrpB to structures present in the Protein Data Bank (PDB).

Our highest resolution structure is an apo structure that diffracted to 1.55 Å. All residues of the native *Cb*PrpB except for four C-terminal residues modelled, including the catalytic loop (Supplementary Table 1). The catalytic loop is designated as residues 113-125 in *Cb*PrpB (Figure 4), harbouring the putative catalytic cysteine. Whilst the catalytic loop region is less well ordered, as in previous structures, we were able to build structure into the region (Figure 5). Between the nine deposited structures, *Cb*PrpB is the most dissimilar structurally (Figure 4), reflecting *C. burnetii’s* position as an evolutionary outlier and possibly correlating with the relatively lower *K_M_* we observe.

**Figure 4:**
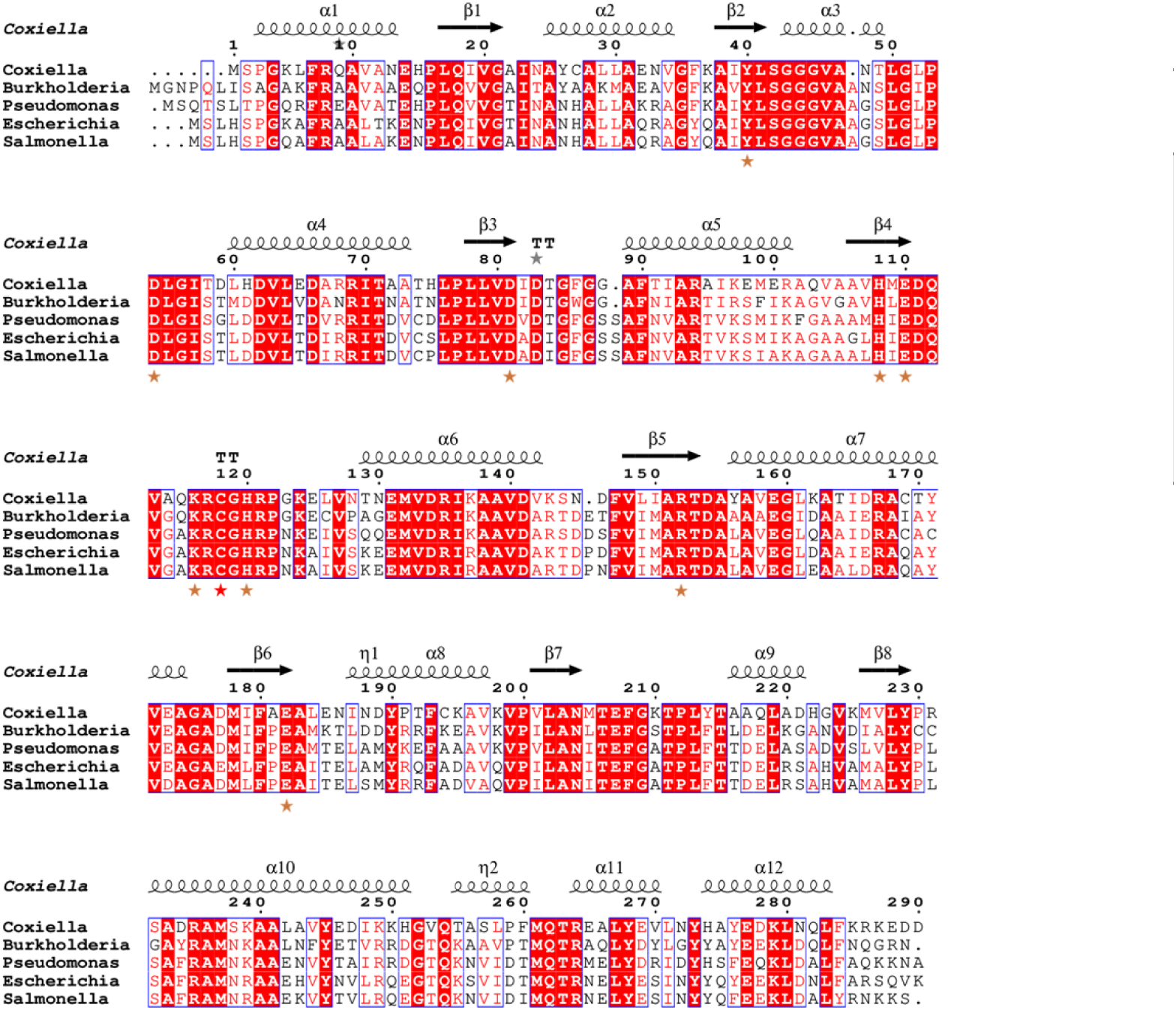
Structural and sequence similarity for selected PrpB structures deposited in the protein databank against C. burnetii. **a**: Calculated C-α root mean square deviation of PDB structures. **b**: Aligned amino acid sequences calculated in ESPript 3 [1] with secondary structure regions displayed. General amino acids discussed here are highlighted with brown stars while cysteine 118 is highlighted with a red star.

**Figure 5:**
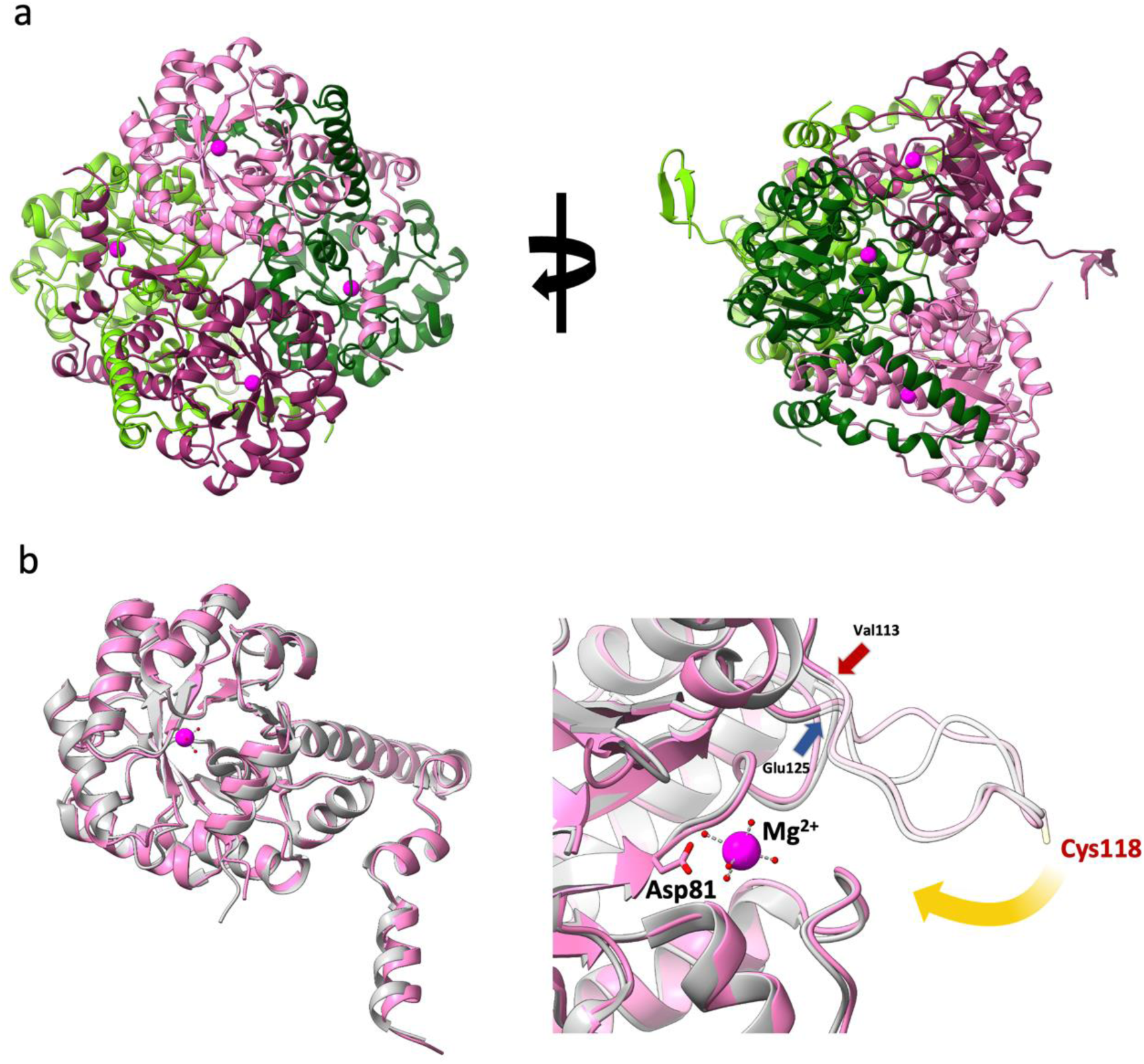
Structural overview for Coxiella burnetii 2-methylisocitrate lyase. Active site magnesium ions are coloured in magenta. **a**: Overview of C. burnetii PrpB tetramer, related by a 90° rotation in the Y axis. **b**: Comparison of a single C. burnetii PrpB protomer (pink) against E. coli (grey, PDB accession 1MUM). Left panel shows overview. Right panel shows close up of active site with start and end catalytic loop residues and cysteine 118 indicated. Key magnesium binding residue Asp81 also indicated. Yellow arrow indicates movement of catalytic loop. All panels generated with UCSF ChimeraX v1.7 [3].

Liu et al. [31] previously published structures of *Escherichia coli* PrpB (*Ec*PrpB) bound to reaction products pyruvate and succinate and the inhibitor isocitrate. These revealed that *Ec*PrpB undergoes a conformational change, with the previously more disordered catalytic loop moving inwards forming a stable, solvent excluded region for lysis to occur. These structures were determined with the expected catalytic cysteine (C118 in *Cb*PrpB) mutated to a serine. Surprisingly, we were able to obtain high resolution complex structures with wild-type *Cb*PrpB (Supplementary Figure 3). These allowed us to observe for the first time an unmodified catalytic loop in the active conformation in a methylisocitrate lyase (Figure 6). We first determined the structure of product-bound *Cb*PrpB (Figure 5). The structure is very similar to that reported for *Ec*PrpB. Isocitrate is bound in a similar conformation between the mutant and wild-type structures. In line with this, residues contacting the products are conserved in *Ec*PrpB and *Cb*PrpB, except for Arg231 substituting for leucine in *E. coli*. On the active site loop, the cysteine sulfur is located within 0.6 Å of the equivalent serine oxygen in the *E. coli* structure. However, the carboxyl group of C5 is moved between the two structures, being around 1 Å further from the cysteine than the serine.

**Figure 6:**
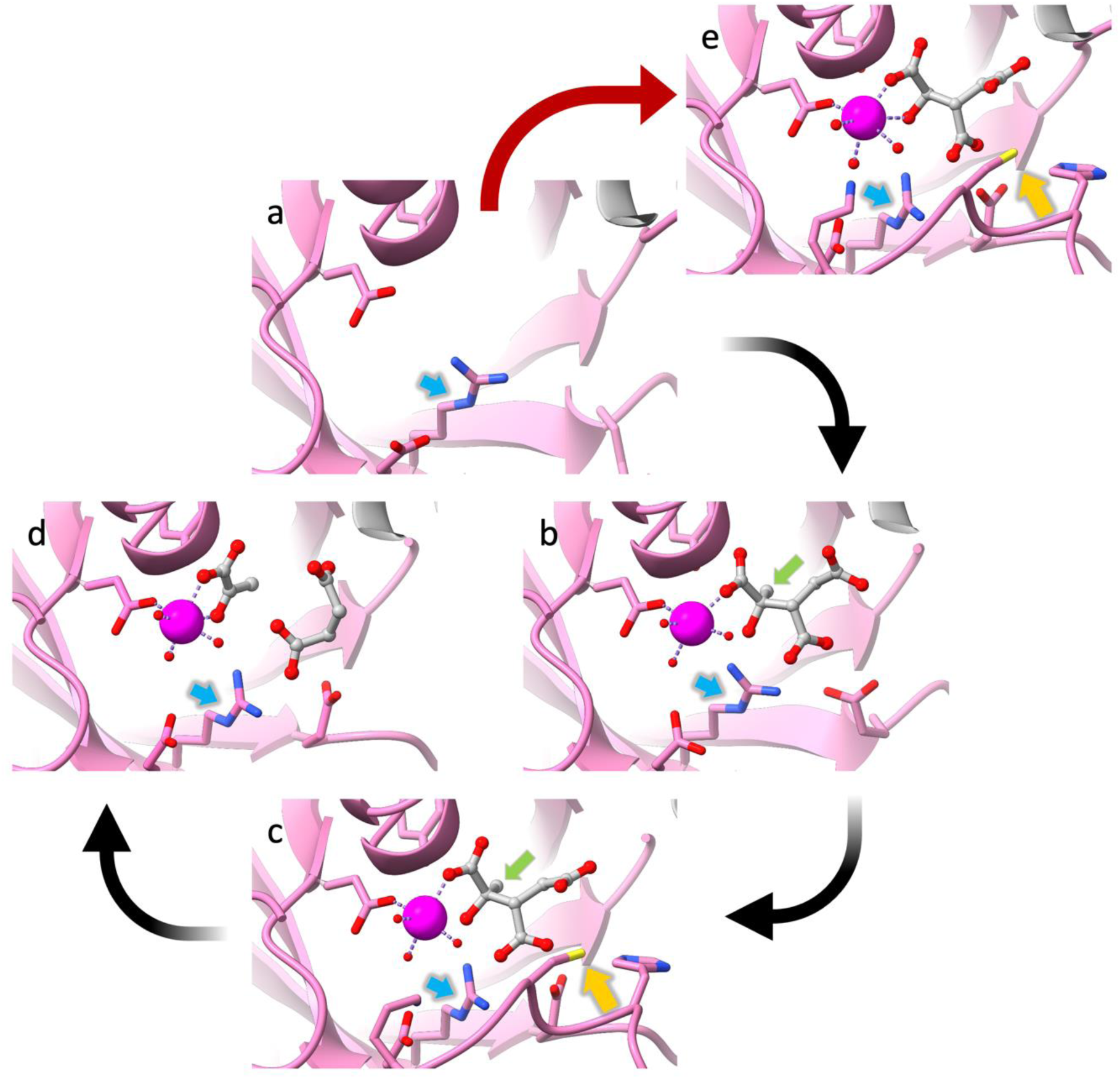
Complete structural analysis of the catalytic pathway for C. burnetii PrpB. Magnesium indicated by magenta atom. Protein coloured in pink, ligand molecules shown in grey stick and ball representation. Oxygen atoms shown in red and nitrogen atoms in blue. **a**: Empty apo active site. **b**: 2-methylisocitrate bound, without loop closure, methyl group indicated by green arrow. **c:** 2-methylisocitrate bound, with loop closure to form the catalytic pose. Catalytic cysteine indicated by yellow arrow. Note how the substrate and R152 subtly reorient upon loop closure to form the pre-reaction state. **d**: products pyruvate and succinate bound. **e**: competitive inhibitor isocitric acid bound. Images produced using UCSF ChimeraX v1.7 [3].

**Figure 7:**
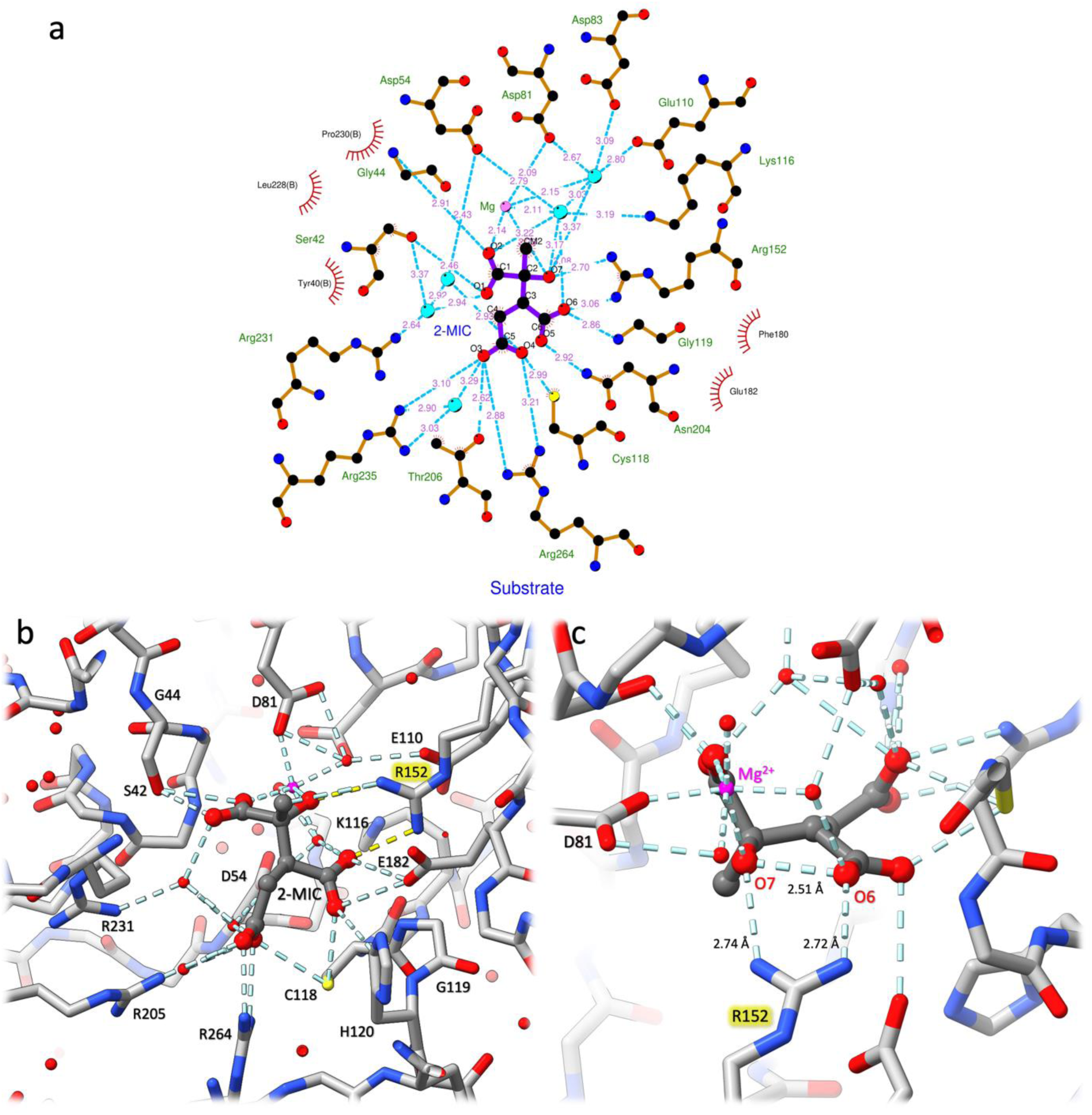
Details of PrpB substrate binding. **a**: 2-D representation of interaction network of the PrpB active site with relevant side chains (brown) and water molecules (cyan). 2-MIC shown in purple and magnesium in pink. **b**: 3-D view of 2- MIC bound matching panel a. Protein shown in grey, 2-MIC shown in dark grey, magnesium ion and waters coloured magenta and red respectively. R152 hydrogen bonds to 2-MIC represented in yellow for clarity. **c**: Close up view focusing on arginine 152. Distance of R152 guanidinium group place it as a likely candidate for the catalytic base, with an intramolecular proton transfer to 2_MIC O6 being a second possibility. Colouring same as panel b. Panel a generated with LigPlot+ v2.2 [2], b and c generated with ChimeraX v1.7 [3].

Next, we determined the structure, at 2.3 Å resolution, of wild-type *Cb*PrpB in complex with the inhibitor isocitrate (Supplementary Tables 1 and 2, Figure 6B). Comparisons with the corresponding *Ec*PrpB complex demonstrates that the two complexes are very similar (RMSD 0.443 Å for superimposition using protein residues less than 5 Å from isocitrate). As with previous structures, isocitrate binding was associated with active site closure and a conformational change in the C-terminus.

To capture a substrate bound structure, crystals were rapidly passed through cryoprotectant containing 5 mM DL-*threo*-2-methylisocitrate and frozen in liquid nitrogen. This yielded the first substrate bound structure of PrpB, at a resolution of 1.9 Å (Figure 6, 7, Supplementary Figure 3, and Supplementary Table 1). Our 2-MIC bound structure showed the substrate binding in a comparable pose to isocitrate in ours and others’ structures. In the substrate bound structure, the two subunits in the crystallographic asymmetric unit occupy different conformations. One protomer displays the active site loop in the active, closed, state whilst the other is in the open, inactive state (Figure 6). Density for the 2-MIC molecule in the inactive protomer was challenging to interpret, indicating either unstable binding or a crystal packing artefact. The protomer with the closed state shows readily interpretably density, with the methyl group clearly visible. We determined a separate isocitrate bound structure showing the closed state: apart from the missing methyl group, the pose of isocitrate is very similar to 2-MIC.

### Structure based inferences on the catalytic mechanism

Upon substrate binding, K116 moves 10.7 Å towards 2-MIC when compared to the inhibitor bound structure, indicating a potential role in catalysis. Indeed, this was one of the previously posited catalytic bases to act as electron donor for the reaction. Moreover, we observe a reorientation of R152 towards the substrate upon binding. R152 particularly approaches a hydrogen bond with the oxygen that is deprotonated to initiate the reaction (Figure 1b) and is positioned perfectly to abstract a proton (Figure 6b).

Building upon our structural data and the hypotheses put forward by Liu *et al*., we designed nine mutants that would enable us to narrow down the catalytic roles of individual residues in the lysis of 2-MIC (Table 2). We tested the catalytic activity of these nine mutants. In agreement with previous studies, mutation of the proposed catalytic acid cysteine to serine (C118S) abrogates all catalytic activity (Table 2, Supplementary Figure 4). Similarly, mutations to Asp54 and Asp81 (D54N, D81N), which coordinate the catalytic magnesium

ion, abolished all activity. Tyr40 and His120 mutants (Y40F and H120Q) showed a large decrease in activity compared to wild-type *Cb*PrpB but remained able to catalyse 2-MIC lysis. These showed *k_cat_* values of 0.6 s^-1^ and 5.7 s^-1^, and *K_M_* values of 0.1 and 4.9 mM, respectively (Table 2).

**Table 2:** Enzyme kinetics of C. burnetii PrpB and active site mutants. All enzymes were tested in a continuous coupled assay using lactate dehydrogenase to reduce the product pyruvate. Rates were detected by fluorescence of the NADH cofactor. The limit of detection was a k_cat_ of 0.06 s^-1^. Errors indicate standard error in the mean from fitting of data in Graphpad v.12 with symmetric standard errors.

| Mutant | $K_M$ (mM) | $k_{cat}$ ($\text{s}^{-1}$) | Role |
| --- | --- | --- | --- |
| WT | 0.39 | ~30,000 | - |
| Y40F | 0.10 | 0.6 | Option for proton shuffle |
| D54N | No Activity | No Activity | Coordinates magnesium |
| D81N | No Activity | No Activity | Coordinates magnesium |
| E110Q | No Activity | No Activity | Option for catalytic base |
| K116Q | No Activity | No Activity | Coordinates magnesium |
| C118S | No Activity | No Activity | Proton donor for reaction |
| H120Q | 4.9 ± 0.8 | 5.7 | Coordinates waters |
| R152Q | No Activity | No Activity | Option for catalytic base |
| E182Q | No Activity | No Activity | Option for catalytic base |

Mutations of Glu110, Lys116, Arg152, and Glu182 (E110Q; K116Q; R152Q; E182Q) also abolished measurable catalytic activity. While Glu182 and Lys116 likely support the positioning of 2-MIC for catalysis, Glu110 and Arg152 are potential catalytic bases.

### Structural analysis of selected mutant structures

E110Q and R152Q were chosen for structural analysis. Both mutants could be crystallised, and structure determination showed that the overall structure was unchanged and that the expected side chain alterations were present (Supplementary Figure 5). Soaking in a large excess (7.5 mM) of substrate yielded only apo structures (Supplementary Figures 5 and 6). Detailed analysis of the active sites revealed that both mutants caused unexpected changes in the hydrogen bonding network of the active site. In the E110Q mutant, the glutamine sidechain moves relative to E110: it no longer forms a hydrogen bond through water with H108 but instead makes a hydrogen bond to D81. This likely makes ligand binding unfavourable. In contrast, in the R152Q mutant, the glutamine sidechain moves to make a new hydrogen bond to H108, displacing the water connecting to E110. This causes a rotation in the pyridine ring of H108 that also requires a rotation in D81. This rotation of D81 disrupts the magnesium coordination pocket, causing the R152Q mutant to no longer bind magnesium or substrate. These structures show that the tight and feature rich active site of PrpB makes dissecting the role of each amino acid highly challenging.

### Discussion

The zoonotic pathogen *C. burnetii* represents both an ongoing human health burden and an economic challenge for livestock owners. The existing therapies require long treatment periods, with the frontline treatment doxycycline causing significant side effects. There is a clear need to identify new antimicrobial targets.

### Incorporating structural information into a downselection process from TraDIS

The TraDIS approach has been used to identify “essential” genes in a wide range of bacteria [45–48]. These studies provide rich datasets but introduce the challenge of down-selecting from many hundreds of essential genes. A common approach has been to compare essential genes between species to identify either common or unique genes. Here, we extended this approach by incorporating structural and druggability prediction. The latest AI enabled tools for protein structure prediction (AlphaFold [41], RosettaFold [49], and ESM-Fold [50]) facilitate robust structure prediction at low cost. AlphaFold version (v2.1.1) was used for this work, as proteins had been selected with a homologue in the PDB. It is likely that there would be limited improvements with AlphaFold 3 [51]. The availability of accurate structure predictions allowed us to assess the druggability of targets. This is an important criterion, as targets unlikely to be amenable to binding of small molecule drugs can be excluded early from the costly development of novel antimicrobials. This approach can significantly de-risk target selection from TraDIS libraries, utilising freely available tools in combination with the code generated in our studies to streamline the process.

### Mechanistic Insights into PrpB Catalysis

As expected, the overall structure of *C. burnetii* PrpB is very similar to that of the *E. coli* ortholog with an RMSD of 0.68 Å and 63% sequence identity. However, kinetic analysis reveals that it is approximately an order of magnitude less active than the *E. coli* enzyme. We find that inhibitor and products bind in a very similar position to that observed for *E. coli*. Remarkably, we have been able to capture the bound substrate. In this structure, the mobile loop closes over the substrate, excluding bulk water but enclosing some water molecules that are candidates for assisting catalysis. Our structures corroborate the observations of Liu et al. (2005) with a similar conformation of the active site loop for *C b*PrpB. The binding of our substrate structure aligns closely to both the inhibitor bound structure presented here and previously. This contradicts the substrate bound model proposed for the *E. coli* enzyme [31] which suggested the substrate would move 1 Å to accommodate the methyl group and approximate the pyruvate binding pose (towards the magnesium). We instead see a significant increase in coordination bond length between the magnesium and the pyruvate moiety. It is possible that the crystal contacts prevent the full evolution of the Michaelis complex and subsequent bond lysis, or that the low pH of our crystals renders the catalytic base unable to perform this role.

The mechanism of the isocitrate and methylisocitrate lyases is generally agreed to start with deprotonation of the hydroxyl group (O2). This forms a carbon-oxygen bond between O2 and C2. This triggers lysis of the C2-C3 bond, forming the first product (glyoxylate or pyruvate respectively). C3 is protonated to form the second product (succinate). Whilst the proton donor is agreed as the catalytic group cysteine, the catalytic base that initiates the reaction remains unclear. Previous suggestions include E110, Y40, and water molecules coordinating the active site magnesium (as proton shuttles), with R152 considered an unlikely alternative [31].

We constructed a battery of structure-guided mutants to dissect the role of specific residues in catalysis. The mutation of Tyr40 to a phenylalanine should remove the ability of tyrosine to act as the catalytic base. The significant residual activity of the Y40F mutant rules out this previously suggested hypothesis for the catalytic base. Two other candidates for the catalytic base, E110Q and R152Q, showed no activity. We crystallised these mutants, revealing no significant changes beyond the mutated residues. However, despite soaking in a high concentration, no substrate binding could not be observed. This indicates a significant decrease in binding affinity for the substrate. Previous investigations into the *M. tuberculosis* isocitrate lyase identified that mutation to alanine of the corresponding residues for E110 and R152 also abolished function [30].

We analysed the position of the water molecules coordinated to magnesium in the substrate structure. These would have their lone pairs orientated away from the substrate hydroxyl. Consequently, we suggest that these are unlikely to perform the proton abstraction that initiates catalysis. A more convincing candidate is R152. The distance from the guanidinium group to O7 of 2-MIC is 2.8 Å, which is favourable for accepting a proton. The movement of R152 observed upon substrate binding also supports this proposal. The major issue with this hypothesis is the very basic *p*K_a_ of arginine, which would normally be protonated at pH 7. However, there may be a local perturbation of the *p*K_a_ by the environment within the active site which could reduce the R152 *p*Ka. Further investigation such as quantum mechanical modelling or neutron diffraction would be needed to corroborate the role of R152.

We also note the close distance of 2.51 Å between the O6 and O7 of 2-MIC when bound within the active site. This is likely indicative of strain within the molecule. This may provide a route for an internal proton shuttle mechanism facilitated by the enzyme-substrate complex, particularly interactions of O6 and O7 with R152.

### Potential of PrpB as a target for antimicrobials against Coxiella burnetii

The structure of the Michaelis complex of *Cb*PrpB with its substrate 2-MIC and our mutants highlight the attractiveness of *Cb*PrpB as a target for antimicrobial development against *C. burnetii*. The structure of the complex demonstrates how the enzyme closes tightly around the substrate, providing a large target pocket as predicted in our downselection cascade. 2- MIC makes numerous interactions with the enzyme that are more extensive than those seen even in ICL, with an additional hydrogen bond between R152 and the 2-MIC hydroxyl formed due to the 2-MIC methyl group driving a rotation in carbon 2 of the substrate compared with isocitrate. The feature rich active site with many potential hydrogen bond donors and acceptors, a metal ion to facilitate compound binding, and bound waters that can provide entropic gains by displacement make *Cb*PrpB attractive for small molecule design. However, a consequence of the intricate network of interactions is that it may be difficult to predict the effect of modest changes, exemplified by the structure of the mutant enzymes, where the tight and feature rich active site of PrpB make dissecting the role of each amino acid highly challenging.

It may also be the case that the relatively high *K_M_* observed for 2-MIC provides an additional benefit for active site drug design as the affinity requirements will be lower for a potential competing compound. PrpB is not essential in several other important bacteria as assessed by TraDIS; this may be because the methylcitrate pathway is not essential in nutrient-rich media, commonly used in TraDIS experiments, whilst *C. burnetii* can only be grown in specialist media designed to simulate its intracellular niche. Alternatively, in many species isocitrate lyase also has activity against 2-MIC. The methylcitrate pathway is essential in *P. aeruginosa* in simulated infection conditions, highlighting that TraDIS can miss useful targets; *Pa*PrpB is essential in these conditions, even though this organism has an isocitrate lyase [43, 52], suggesting that an inhibitor of *Cb*PrpB might be efficacious in a broader range of species.

In conclusion, our structure of *Cb*PrpB in complex with its substrate 2-MIC suggests that the catalytic base for methylisocitrate lyase is R152. Our structures of mutants of prospective bases highlight that the (methyl)isocitrate lyase active site is highly coordinated, making the development of resistant mutations that retain catalytic activity less likely. As an essential protein that is readily druggable, PrpB offers an attractive opportunity for development of novel antimicrobials targeting *C. burnetii*.

## Materials and Methods

### Target Curation, Protein Structure Prediction and Pocket Scoring

Essential gene data was incorporated into a Microsoft Excel spreadsheet for scoring. AlphaFold v2.1.1 was installed utilising the non-Docker install from Kalininalab on GitHub (https://github.com/kalininalab/alphafold_non_docker) on a workstation running Ubuntu 20.04.3 with an AMD Ryzen Threadripper PRO 3975X CPU and one Nvidia RTX 3090 GPU (used for all computational tasks). AlphaFold predictions were run with a cut-off date for PDB templates of 10^th^ November 2021. Pocket location and scoring utilised P2RANK 2.4 running with the default configuration file [42]. The automated script for assembling AlphaFold inputs is available on GitHub.

### CbPrpB Purification

The wild-type *C. burnetii* RSA493 PrpB (CBU_0771; Q83DG5) was codon optimised for *E. coli* using an in-house script [53]. The optimised gene and point mutants were synthesised and inserted in pNIC28-Bsa4 [54] and ordered from Twist Biosciences. All constructs were transformed into *E. coli* BL21 (DE3) cells (Merck #69450-3) alongside the pGro7 chaperone plasmid (Takara #3340). Cells were grown at 37°C, 220 RPM in LB media supplemented with kanamycin (50 µg/mL) and chloramphenicol (25 µg/mL) until OD_600_ reached 0.6. Cells were then treated with 3 mg/mL L-arabinose (Melford, #A51000-100.0) to induce expression of chaperones and incubation temperature reduced to 16 °C. After 30 minutes, *Cb*PrpB expression was induced with 200 µM isopropylthio-β-D-galactoside (BioSynth, #EI07011) and cells harvested after 48 hours. Cells were resuspended in 20 mM Tris-HCl, 500 mM NaCl, pH 8.0 (Buffer A). Cells were lysed with sonication using an Ultrasonics VCX-130 on ice with an amplitude of 70%, 8 s on/off for 8 minutes. Cell lysate was centrifuged at 22,000 x*g* for 45 minutes at 4 °C and the supernatant filtered to 0.22 µM. Clarified lysate was added to ∼ 1 mL nickel-sepharose 6 fast flow resin (Ni-sepharose 6 fast flow, Cytiva #10249123) and rolled at 4 °C for 45 minutes. Beads were isolated through gravity flow (Econo-Pac, Bio-Rad #7321010) and washed with Buffer A supplemented with 20 mM imidazole pH 8.0. *Cb*PrpB was eluted in Buffer A supplemented with 500 mM imidazole pH 8.0. *Cb*PrpB was polished on a Superdex 200 16/600 size-exclusion chromatography column (Cytiva #28989335) using an Äkta Pure 25 L (Cytiva #29018224) and eluted isocratically in 20 mM HEPES-NaOH, 200 mM NaCl, 1 mM dithiothreitol, pH 7.5. Fractions containing purified *Cb*PrpB were collected and concentrated to working concentrations with a regenerated cellulose 30 kDa Amicon Ultra 15 mL (Merck, #UFC903008) and 10 kDa Amicon Ultra 0.5 mL (Merck, #UFC503008).

The final product was over 95% pure by SDS-PAGE.

Protein for crystallisation was stably concentrated to up to ∼60 mg/mL, snap frozen in liquid nitrogen and stored at -80 °C. Protein for kinetic analysis was diluted to 10 nM in the size exclusion buffer supplemented with 25% glycerol (v/v) and stored at -20 °C. Protein remained active after many freeze thaw cycles in the glycerol supplemented buffer.

Mutants for assays and crystallisation were prepared using the same protocol as wild-type.

### CbPrpB Crystallisation

Wild-type crystals not soaked with substrate were set up using an Oryx 8 crystallisation robot (Douglas Instruments). Oryx grown crystal plates were set up with *Cb*PrpB at a final concentration of 10 mg/mL mixed in equal parts with mother liquor (10% v/v ethanol, 1% w/v PEG 1000, 0.1 M sodium-citrate buffer pH 4.2) and grown under oil in microbatch format. Substrate bound and mutant crystals were grown using a Mosquito crystallisation robot (SPT LabTech). Mosquito grown crystals were grown in sitting drop vapour diffusion using MRC 96-Well-2-Drop crystallisation plates (Molecular Dimensions #MD11-00-40) with 100 or 200 nL drop sizes and 60 µL reservoir. E110Q and R152Q mutant crystals were prepared using the Mosquito.

Apo crystals were grown in mother liquor optimised in-house from condition JCSG+ crystal screen condition B6 (Molecular Dimensions, #MD1-40). Crystals used for substrate bound structures were grown in; 40% v/v ethanol, 5% w/v PEG 1000, 0.1 M sodium-citrate buffer pH 4.2. Isocitrate bound structures were grown in; 0.3 M sodium acetate trihydrate, 0.1 M sodium cacodylate pH 6.5, 8 % w/w PEG 20 000 and 8% v/v PEG 500 MME. Crystals for the product bound structure were grown in; 0.1 M MES pH 6.0, 0.2 M NaCl, 20% w/w PEG 6000 and 10% v/v ethylene glycol.

All *P*3_1_21 crystals (apo, substrate bound and mutants) were frozen in their mother liquor supplemented with 30% v/v DMSO. The wild-type substrate bound structure cryo- protectant contained 5 mM 2-MIC, increased to 7.5 mM for both mutant crystals. Isocitrate bound crystals were frozen in mother liquor supplemented with 30% DMSO and 20 mM isocitrate. Product bound crystals were frozen in 30% v/v DMSO, 20% w/w PEG 3350, 100 mM NaCl, 40 mM pyruvate and 40 mM succinate.

### Structure Determination

Frozen crystals were sent to Diamond Light Source in Oxfordshire, UK for unattended data collection on beamlines i03, i04 and i24. Xia2 [55] reduced data from Diamond in-house autoprocessing were taken for further processing. Molecular replacement was carried out with Phaser [56] utilising the AlphaFold 2.3 predicted structure of *Cb*PrpB [41]. Refinement was carried out using Refmac5 [57] in CCP4 Cloud [58]. Models were rebuilt in Coot 0.9 [59] and Isolde 1.7 [60] within ChimeraX [3].

### PrpB Assay

DL-*threo*-2-methylisocitrate (Biosynth #WCA18366) was resuspended at 50 mM in water and stored at -20 °C. Reaction mix contained: 5 mM DTT, 2.5 mM MgCl_2_, 50 mM HEPES pH 7.5, 250 µM NADH, 1.5 µg L-LDH (Roche #10127230001) and 50 pM PrpB with 2-MIC and inhibitor if added at appropriate concentrations. Plates were incubated at 37 °C before the reaction and initiated by addition of 2-MIC. The NADH concentration was monitored by fluorescence with excitation and emission wavelengths of 340 and 440 nm, respectively. The limit of detection was defined as two times the error in a no enzyme control (*n* = 6).

## Acknowledgements

The authors thank the staff of Diamond beamlines I03, I04, and I24 for their support in collecting protein structure data.

