## Supplementary information for "Structure and catalytic mechanism of methylisocitrate lyase, a potential drug target against Coxiella burnetii"

**Supplementary Figure 1**: **Purification of recombinant *C. burnetii* PrpB from *E. coli***. PrpB was co-expressed with the chaperone GroEL/ES in order to yield soluble protein. Gravity flow nickel affinity chromatography was used to isolate PrpB from crude cell lysate using an N-terminal 6x histidine tag. Partially purified protein was concentrated and polished using a Superdex 200 pg 16/600 column (Cytiva #28989335) on an ÄKTA Pure system. **a**: 4-20% SDS-PAGE (GenScript #M42015) using 10 µL gel filtration sample mixed 1:1 with SDS loading buffer and heated at 90 °C for 5 minutes. Marker used is Spectra Multicolour Broad Range Protein Ladder (Thermo Scientific #26634). Gel was imaged using a BioRad ChemiDoc XRS+ (#170-8265). Denatured PrpB subunits run between 30 and 35 kDa (red box). **b**: Chromatogram of PrpB gel filtration from ÄKTA Pure. Peak for PrpB indicated by red line. Data from Unicorn 7.9 (Cytiva) imported into GraphPad Prism 10 for display.

a

b

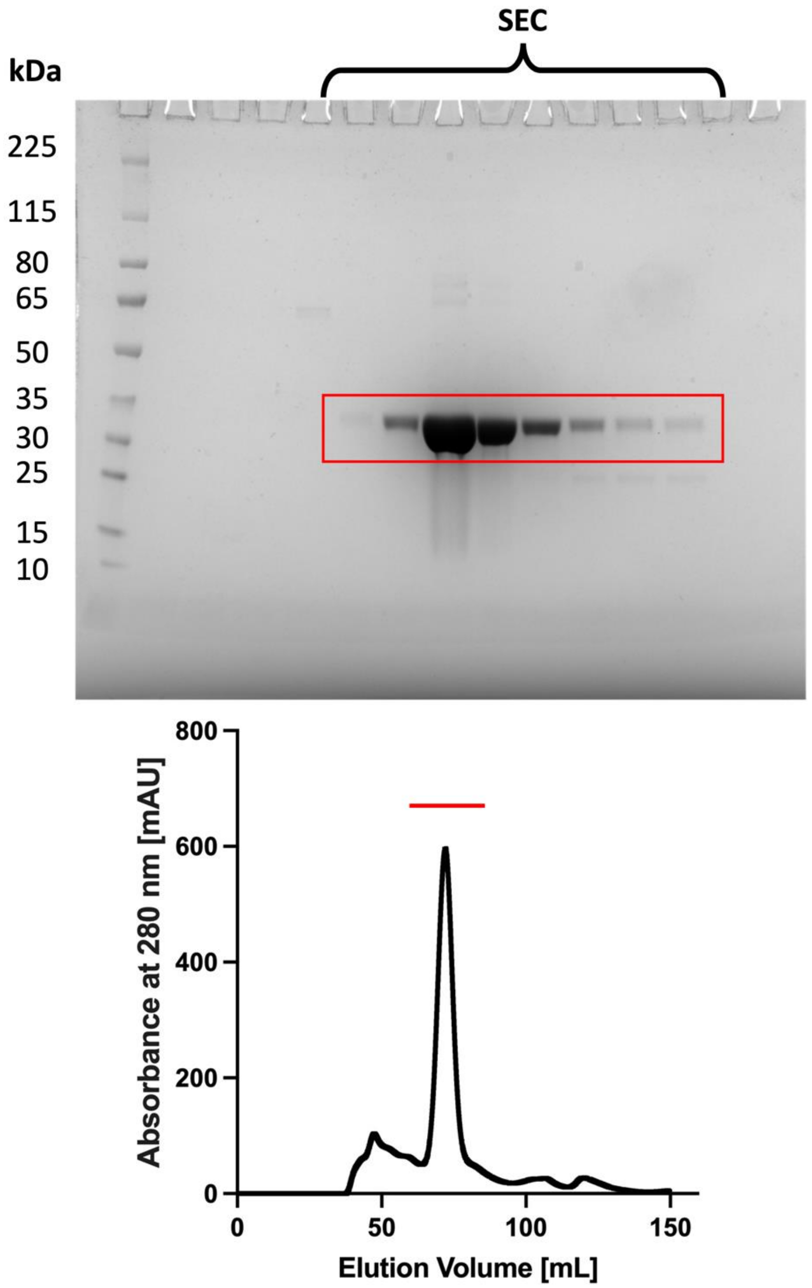

**Supplementary Figure 2**: **Differential scanning fluorimetry analysis of *C. burnetii* PrpB supports a multimeric structure**. Sample was tested at a concentration of 0.1 mg/mL with 8X SYPRO Orange (Invitrogen #S6650) in 10 mM Hepes pH 7.0, 150 mM NaCl. Differential scanning fluorimetry was performed on a QuantStudio6 qPCR machine (Applied Biosystems), with temperature increasing at a rate of 1 °C per minute, with fluorescence read every 1 °C from 25 to 95 °C. Data were analyzed using Protein Thermal Shift software v1.4 (Applied Biosystems). **a**: Raw data fluorescence data. **b**: Derivative data. Twin peaks observed in the derivative data show a two stage unfolding (green arrows), likely through initial disassembly of the tetramer followed by unfolding of monomeric PrpB.

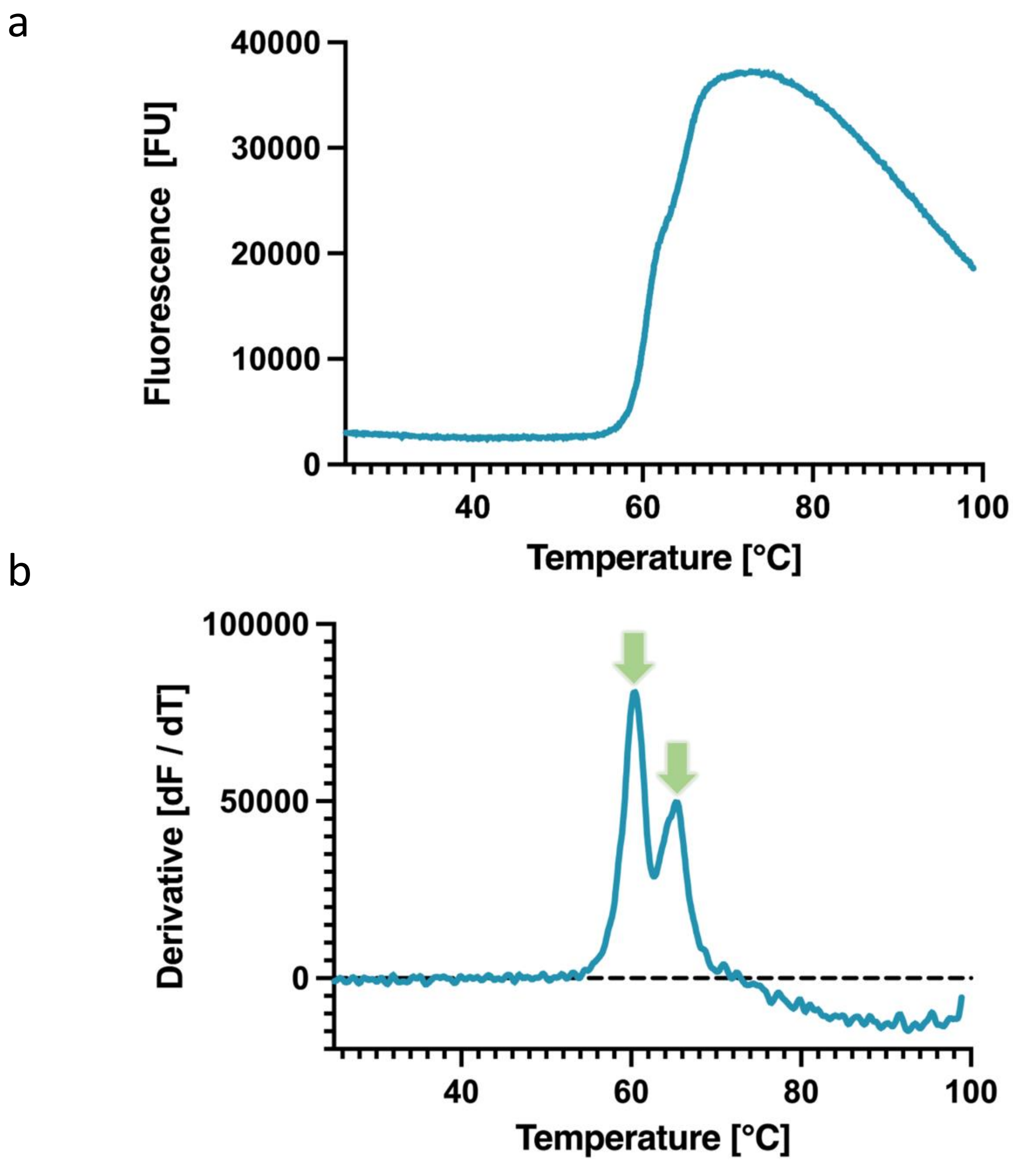

**Supplementary Figure 3: Density for observed for bound wild-type PrpB structures**. **a**: Isocitrate bound to PrpB. As a competitive inhibitor of 2-methylisocitric acid, isocitrate is observed to bind in an essentially identical pose to substrate. **b**: Products from the lysis of 2-methylisocitric acid, pyruvate and succinate bound to PrpB. Pyruvate is seen to coordinate the magnesium while succinate is distant. Product release is likely ordered with succinate release enabling pyruvate to leave. **c** and **d**: 2-methylisocitric acid bound to the wild-type enzyme. Clear density is seen for the unique methyl group (green arrow, panel **d**), as with the sulfur atom of C118. Panel **d** shows a close-up view of pyruvate group.

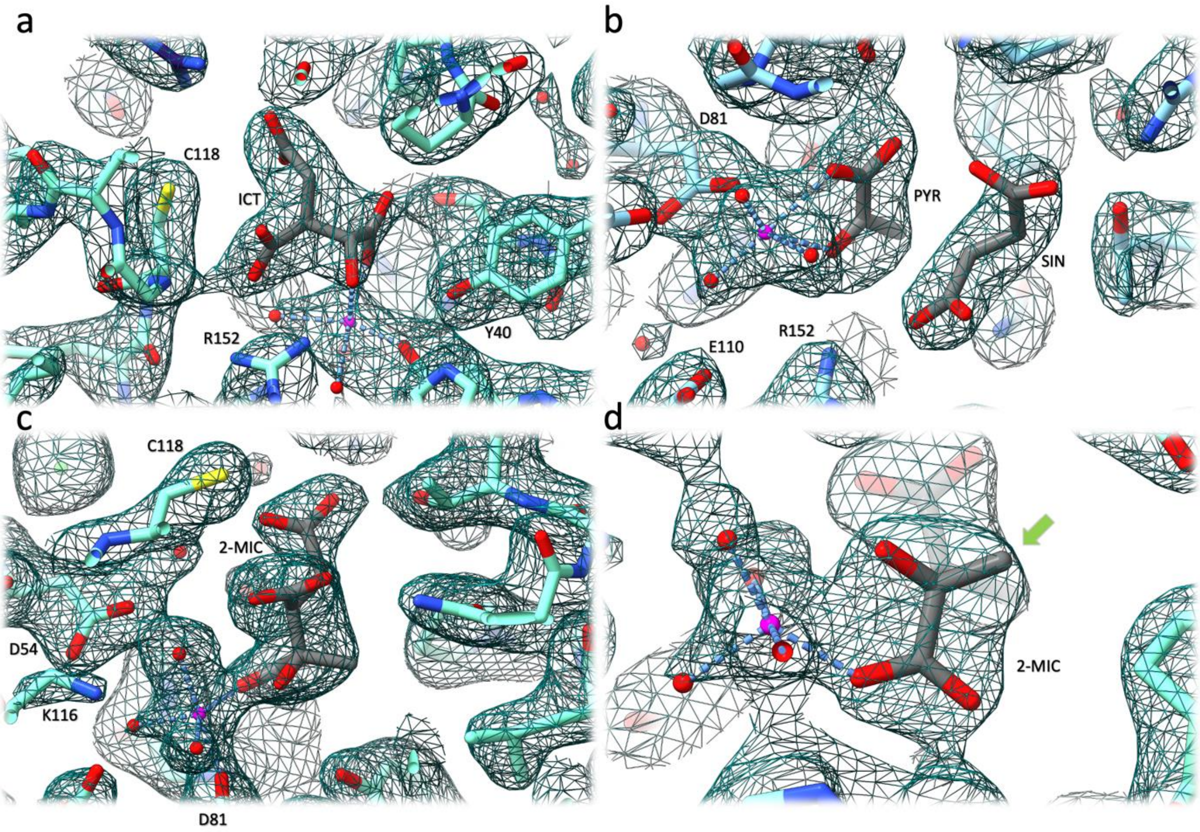

**Supplementary Figure 4: Kinetic analysis of mutant PrpB and limit of detection.** Activity remained in mutants Y40F (**a**) and H120Q (**b**). Experiments to determine kinetic parameters were limited by the maximum concentration of substrate available. Y40F assay was performed as with wild type but at an enzyme concentration of 1.25 µM, allowing determination of kinetic parameters. H120Q assay carried out at an enzyme concentration of 300 nM. The estimate of $K_M$ is less accurate as concentrations significantly above $K_M$ could not be tested. **c**: Raw data from example inactive assay. C118S assay run with a negative (no enzyme) and positive control (wild type at 100 pM). No detectable C118S activity was observed up to 10 µM. Limit of detection of the assay is reached at between 10-20 pM wild-type enzyme, indicating that the inactive mutants are at least $5x10^5$ fold slower than wild-type. All panels generated with GraphPad Prism.

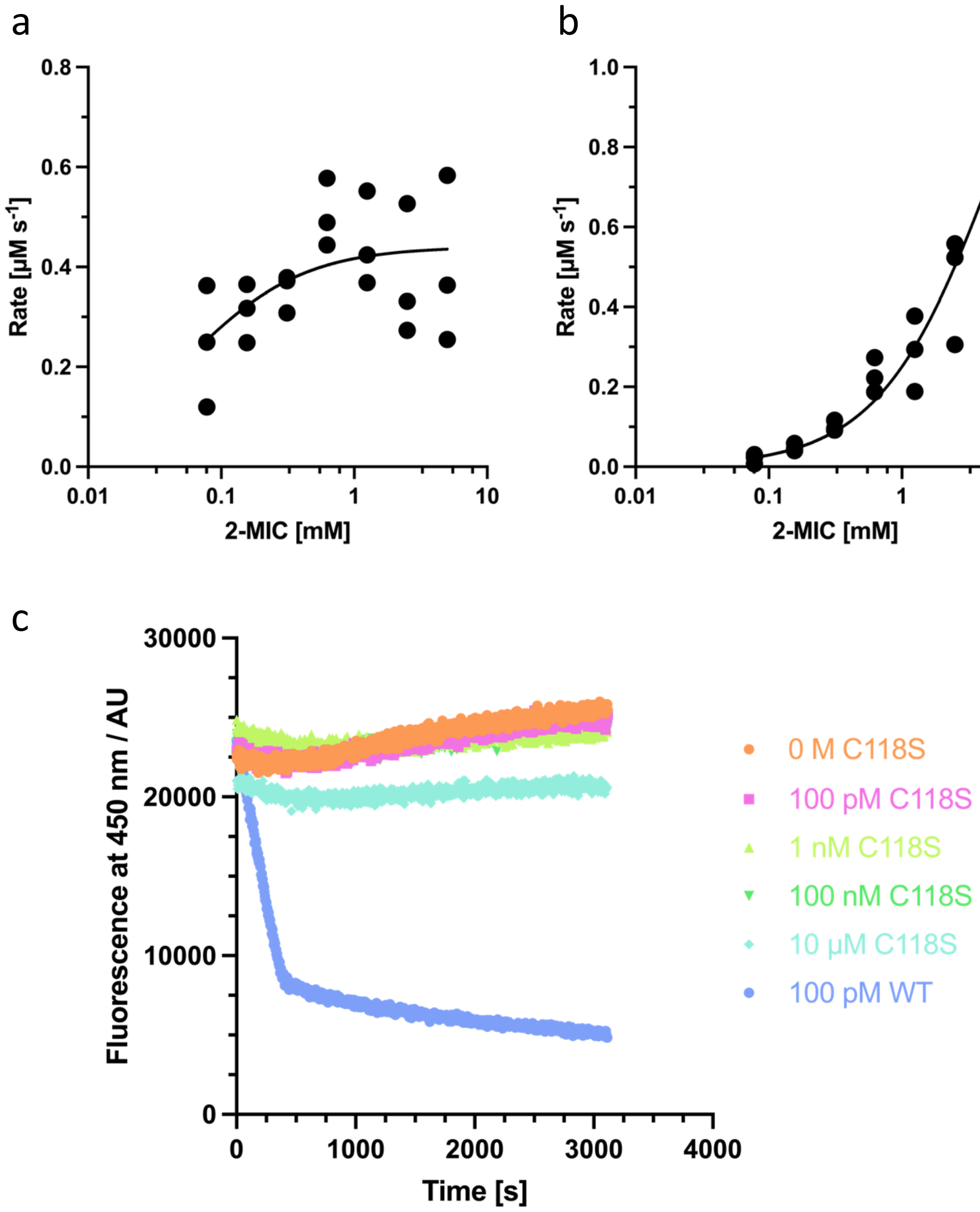

**Supplementary Figure 5: Structural overview of PrpB mutants and corresponding electron density**. For all panels: green represents wild-type, purple represents E110Q and pink R152Q. Oxygen atoms are labelled red and nitrogen atoms blue. Density shown is a $2F_o$-$F_c$ map (contoured at $\sigma$: 0.688 **b** & 0.690 **c**, 1.401 **d** & 1.390 **e**). **a**: Mutant structures aligned against the wild-type PrpB structure. Overall fold remains unaffected showing that loss of

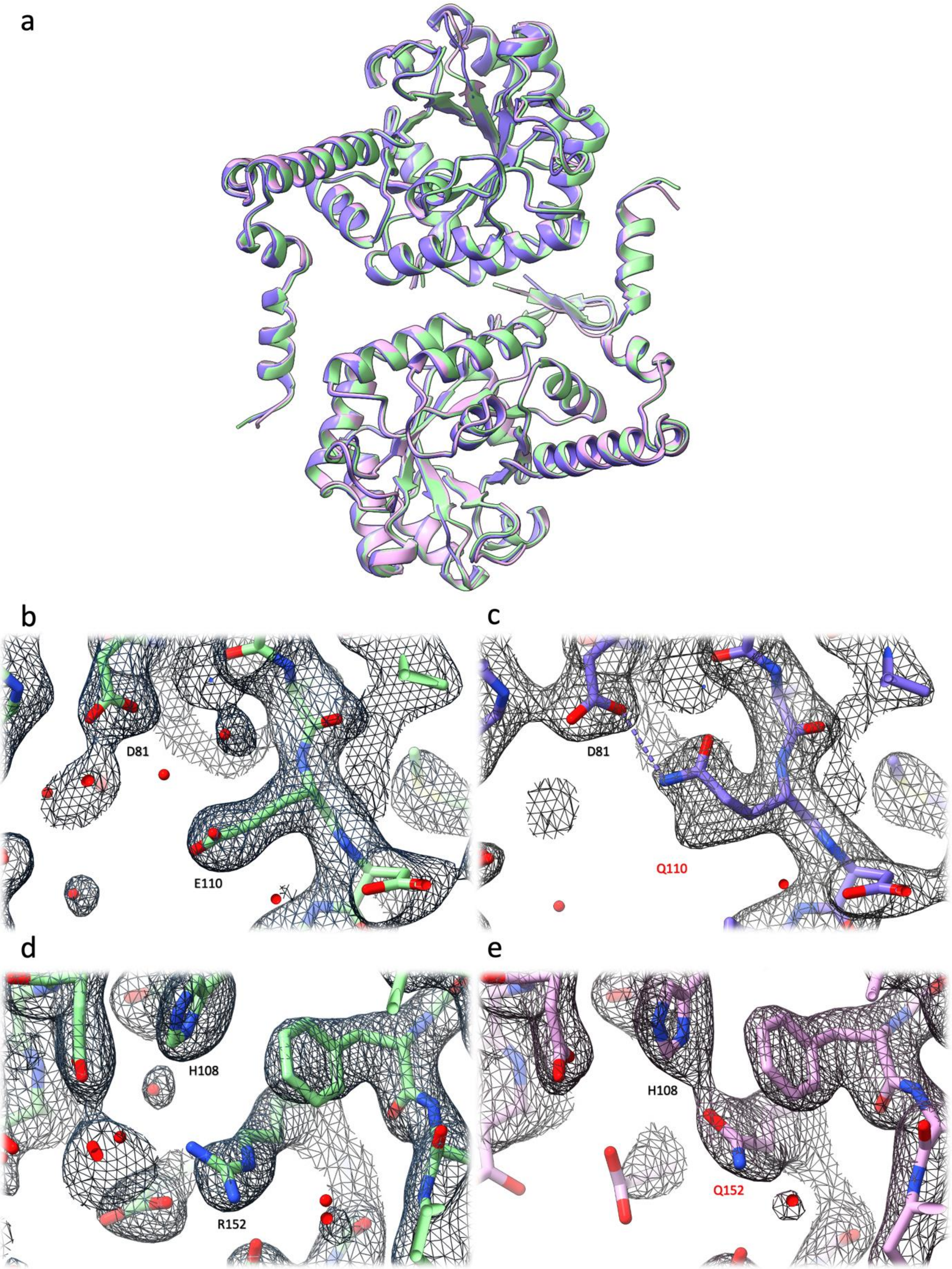

activity is due to mutation and active site disruption rather than protein misfolding. Panels **b** and **c** show effect of the loss of glutamic acid 110 compared to wild-type. Despite the conservative mutation, glutamine adopts a twisted conformation, making a new hydrogen bond to aspartic acid 81. Glutamine replacing R152 induces a new hydrogen bond with H108, in turn causing the loss of a water molecule (panels **d** and **e**). All panels generated with UCSF ChimeraX v1.7.

**Supplementary Figure 6: Dissecting the impact of mutants on active site structure**. Green represents wild-type, purple represents E110Q, and pink R152Q. Oxygen atoms are labelled red and nitrogen atoms blue. Relevant amino acids are labelled. Panel **a** shows wild-type, **b** shows E110Q mutant, **c** shows R152Q and **d** shows all aligned. All views are the same. All panels generated with UCSF ChimeraX v1.7.

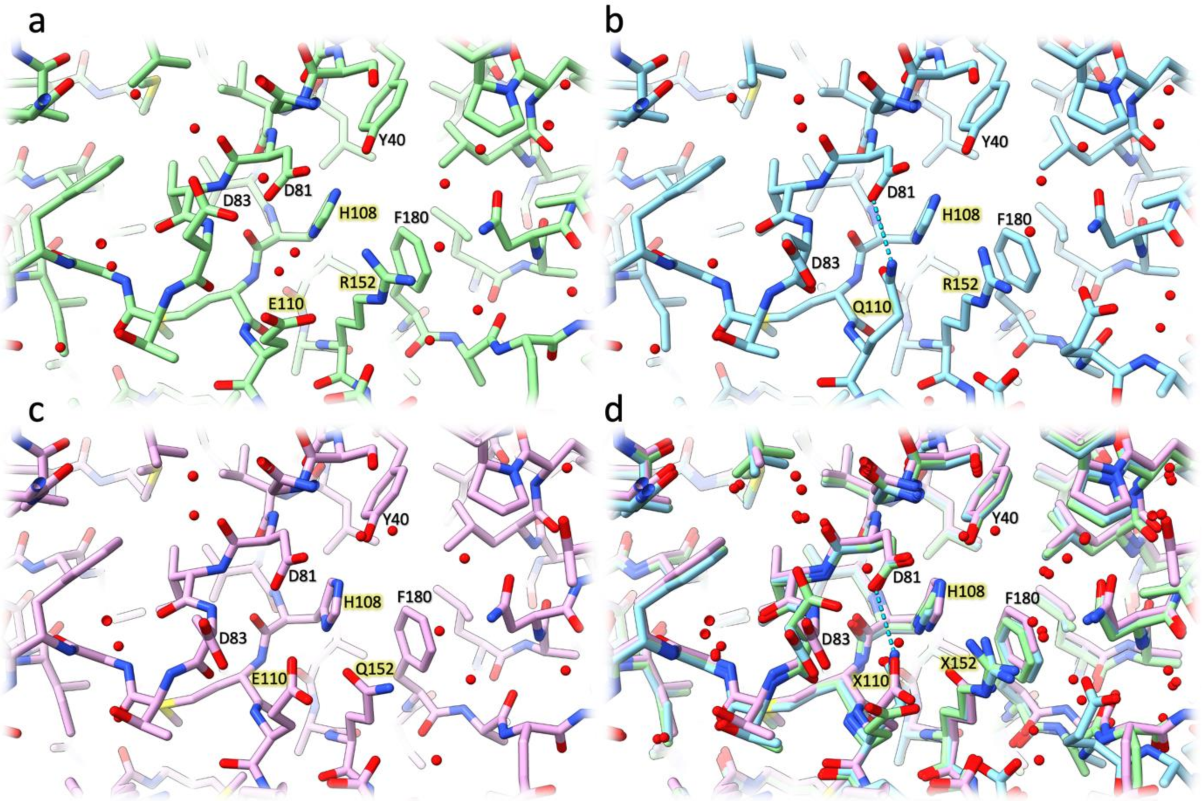

**Supplementary Table 1: Crystallographic table.** Data for the highest resolution shell are shown in parentheses. All data were collected at Diamond Light Source.

| Dataset | Apo | 2-MIC | Product | Isocitrate | E110Q | R152Q |
|---|---|---|---|---|---|---|
| PDB ID | 9HGK | 9HGQ | 9HRA | 9HHY | 9HGO | 9HHS |
| Asymmetric Unit | 2 protomers | 2 protomers | 8 protomers | 12 protomers | 2 protomers | 2 protomers |
| Beamline | I03 | I04 | I04 | I03 | I24 | I24 |
| Wavelength (Å) | 0.9763 | 0.8517 | 0.9537 | 0.9763 | 0.9999 | 0.9999 |
| Resolution range | 44.58 - 1.6 (1.62 - 1.6) | 63.49 - 1.9 (1.94 - 1.9) | 52.59 - 2.48 (2.51 - 2.48) | 54.47 - 2.33 (2.36 - 2.33) | 61.48 - 1.82 (1.86 - 1.82) | 52.58 - 1.7 (1.73 - 1.7) |
| Space group | *P* 31 2 1 | *P* 31 2 1 | *P* 1 21 1 | *P* 1 21 1 | *P* 31 2 1 | *P* 31 2 1 |
| Unit cell | 74.96 74.96 183.99 90 90 120 | 73.31 73.31 184.141 90 90 120 | 105.199 101.203 115.683 90 91.15 90 | 93.785 115.55 195.241 90 92.34 90 | 74.18 74.18 184.45 90 90 120 | 73.94 73.94 184.23 90 90 120 |
| Total reflections | 1,774,681 (89,245) | 948,419 (44,540) | 606,616 (27,108) | 1,252,024 (62,653) | 450,561 (25,757) | 614,001 (31,090) |
| Unique reflections | 77749 (1590) | 45323 (2064) | 85869 (2739) | 177371 (5817) | 51028 (2833) | 65133 (2784) |
| Multiplicity | 20.18 (20.67) | 20.54 (19.68) | 7.05 (6.47) | 7.04 (7.09) | 8.70 (9.87) | 9.42 (9.65) |
| Completeness (%) | 97.28 (56.68) | 98.26 (72.80) | 99.72 (95.70) | 99.83 (98.13) | 95.04 (95.68) | 99.98 (99.96) |
| Mean I/sigma(I) | 17.64 (0.26) | 14.94 (0.42) | 4.83 (0.58) | 8.29 (0.45) | 13.32 (0.50) | 17.62 (0.90) |
| Wilson B-factor | 39.53 | 41.49 | 41.94 | 53.96 | 37.64 | 39.88 |
| R-merge | 0.0735 (6.9279) | 0.1239 (3.8752) | 0.4226 (3.0058) | 0.1676 (2.5944) | 0.1423 (4.2280) | 0.0483 (2.6106) |
| R-meas | | | | | | |
| R-pim | | | | | | |
| CC1/2 | 0.9998 (0.3264) | 0.9996 (0.3986) | 0.9695 (0.2558) | 0.9945 (0.2991) | 0.9952 (0.3443) | 0.9995 (0.3025) |
| Reflections used in refinement | 77749 (1590) | 45323 (2064) | 85869 (2739) | 177371 (5817) | 51028 (2833) | 65133 (2784) |
| Reflections used for R-free | 3855 (64) | 2219 (75) | 4243 (156) | 8837 (299) | 2589 (157) | 3243 (133) |
| R-work | 0.2348 | 0.1888 | 0.2081 | 0.214 | 0.2321 | 0.2104 |
| R-free | 0.2989 | 0.2429 | 0.2513 | 0.2652 | 0.2989 | 0.2584 |
| CC(work) | | | | | | |
| CC(free) | | | | | | |
| Number of non-hydrogen atoms | 4821 | 4753 | 17963 | 27733 | 4771 | 4842 |

| | | | | | | |
|---|---|---|---|---|---|---|
| macromolecules | 4526 | 4394 | 17149 | 26526 | 4499 | 4551 |
| ligands | 40 | 88 | 271 | 326 | 34 | 34 |
| solvent | 255 | 271 | 543 | 881 | 238 | 257 |
| Protein residues | 587 | 572 | 2243 | 3456 | 584 | 587 |
| RMS(bonds) | 0.009 | 0.009 | 0.009 | 0.008 | 0.009 | 0.01 |
| RMS(angles) | 1.43 | 1.47 | 1.42 | 1.46 | 1.58 | 1.52 |
| Ramachandran favored (%) | 97.6 | 97.54 | 98.24 | 98.46 | 97.07 | 97.6 |
| Ramachandran allowed (%) | 2.4 | 2.29 | 1.76 | 1.54 | 2.93 | 2.4 |
| Ramachandran outliers (%) | 0 | 0.18 | 0 | 0 | 0 | 0 |
| Rotamer outliers (%) | 2.77 | 1.32 | 1.75 | 1.79 | 3.43 | 1.9 |
| Clashscore | 4.58 | 3.89 | 4.41 | 3.9 | 5.38 | 4.68 |
| Average B-factor | 57.88 | 50.53 | 53.95 | 68.95 | 55.06 | 55.81 |
| macromolecules | 58.22 | 50.34 | 54.34 | 69.41 | 55.28 | 55.91 |
| ligands | 66.32 | 62.55 | 61.32 | 81.39 | 59.06 | 66.09 |
| solvent | 50.38 | 49.69 | 37.74 | 50.69 | 50.27 | 52.78 |

**Supplementary Table 2: Details of PrpB crystallisation conditions**.

| **Dataset** | **Protein Buffer** | **Mother Liquor** | **Cryoprotectant** |
|---|---|---|---|
| **Apo** | 200 mM NaCl, 20 mM HEPES 7.5 | 10% v/v EtOH, 0.1 M sodium-citrate pH 4.2, 1% w/w PEG 1000 (in house) | 30% v/v DMSO, mother liquor |
| **Substrate** | 200 mM NaCl, 20 mM HEPES 7.5 | 40% v/v EtOH, 0.1 M sodium-citrate pH 4.2, 5% w/w PEG 1000 (JCSG+, B6) | 30% v/v DMSO, mother liquor, 5 mM 2-MIC |
| **Inhibitor** | 200 mM NaCl, 20 mM HEPES 7.5 | 0.3 M sodium acetate trihydrate<br>0.1 M sodium cacodylate 6.5<br>8% w/v PEG 20 000<br>8% v/v PEG 500 MME<br>(Clear Strategy I, D7) | 30% v/v DMSO + 20 mM isocitrate |
| **Product** | 200 mM NaCl, 20 mM HEPES 7.5 | 0.1 M MES pH 6<br>0.2 M NaCl<br>20% w/v PEG 6000<br>10% v/v ethylene glycol<br>(Ligand Friendly Screen, B7) | 30% v/v DMSO, 20% w/w PEG 3350, 100 mM NaCl, 40 mM pyruvate, 40 mM succinate |
| **R152Q** | 200 mM NaCl, 20 mM HEPES 7.5 | 10% v/v EtOH, 0.1 M sodium-citrate pH 4.2, 1% w/w PEG 1000 (in house) | 30% v/v DMSO, mother liquor, 7.5 mM 2-MIC |
| **E110Q** | 200 mM NaCl, 20 mM HEPES 7.5 | 10% v/v EtOH, 0.1 M sodium-citrate pH 4.2, 1% w/w PEG 1000 (in house) | 30% v/v DMSO, mother liquor, 7.5 mM 2-MIC |
